# The virus-encoded ion channel “viroporin” activity of the agnoprotein is required for BK Polyomavirus release from infected kidney cells

**DOI:** 10.1101/2023.01.30.526200

**Authors:** Gemma Swinscoe, Emma L Prescott, Daniel L Hurdiss, Ethan L Morgan, Samuel J Dobson, Thomas Edwards, Richard Foster, Matthew Welberry Smith, Andrew Macdonald

## Abstract

BK polyomavirus (BKPyV) is a common opportunistic pathogen and the causative agent of several diseases in transplant patients and the immunosuppressed. Despite its importance, aspects of the virus lifecycle such as how the virus exits an infected cells, remain poorly understood. The late region of the BKPyV genome encodes an auxillery protein called agnoprotein. We and others have shown that agnoprotein is an essential factor in virus release, and the loss of agnoprotein results in an accumulation of virus particles within the nucleus of an infected cell. The functions of agnoprotein necessary for this egress phenotype are not known. Here we demonstrate that agnoprotein shows properties associated with viroporins, a group of virus-encoded membrane spanning proteins that play key roles in virus infection and release. We demonstrate that agnoprotein oligomerises and perturbs membranes in cells. The development of a novel recombinant agnoprotein expression system permitted the identification of the first small molecules targeting agnoprotein. These compounds abrogated agnoprotein viroporin activity in vitro and reduced virus release, indicating that viroporin activity contributes to the phenotype observed in agnoprotein knockout viruses. The identification of channel activity should enhance the future understanding of the physiological function of agnoprotein and could represent an important target for antiviral intervention.

## Introduction

Polyomaviruses are small, double-stranded DNA (dsDNA) viruses, which infect a range of mammals, birds, and fish (DeCaprio & Garcea, 2013). Fourteen human polyomaviruses (hPyV) have been discovered since their initial discovery in 1971. All hPyVs cause life-long chronic infection in humans with seroprevalences ranging from 10-95% in the adult population, however, only the minority are associated with disease (Kamminga et al. 2018). The clinical relevance of polyomavirus-associated disease is associated with immunocompromised transplant patients, or patients who have acquired/congenital immunodeficiencies (De Gascun & Carr, 2013).

BK polyomavirus (BKPyV) is a major etiological factor of polyomavirus-associated nephropathy (PVAN) and urethral stenosis in renal transplants, and late-onset haemorrhagic cystitis in hematopoietic stem cell transplants. Primary infection with BKPyV is thought to occur during early childhood and leads to chronic subclinical infection. 90% of the adult population is chronically infected with BKPyV, and while approximately 70% of infected individuals experience low level asymptomatic urinary shedding at any given time, healthy immunocompetent individual successfully control virus reactivation (DeCaprio & Garcea, 2013). BKPyV reactivation is more serious if an individual is immunocompromised. Viral reactivation in the absence of a competent immune response often leads to PVAN, and more rarely meningoencephalitis, bilateral atypical retinitis, and interstitial pneumonitis. 10% of renal transplant patients develop PVAN, and 90% of these cases result in acute transplant rejection (Kant et al. 2020; Hirsch et al. 2002; Hirsch et al. 2005; Dharnidharka et al. 2009; Schold et al. 2009). Although, PVAN causes a substantial impact on both health care systems and the patients affected, there are no direct acting antivirals against BKPyV (Johnston et al. 2010). Current treatment guidelines advise lowering a patient’s immunosuppressive therapies, and the general antiviral drug; cidofovir, which is nephrotoxic and rarely prescribed to PVAN patients by clinicians. There is desperate need for the development of antivirals to combat PVAN, and by understanding the lifecycle of BKPyV new antivirals can be better targeted to undermined viral processes.

BKPyV virions contain a ~5 kbp genome, which is divided into three functional regions: early and late encoding regions that are separated by the non-coding control region (NCCR). The NCCR control region contains the viral origin of replication and a bidirectional promoter that serves the early and late encoding regions (Chong et al. 2019). Recombinations in the NCCR are found in polyomavirus-associated diseases and are thought to contribute to the reactivation and pathogenesis of polyomaviruses. The early region encodes for the large tumour antigen (LT), small tumour antigen (sT), and alternatively spliced variants of LT, which are essential for virus transcription and replication (Moens & Macdonald, 2019). The late region encodes the structural proteins: VP1, VP2, VP3, and a poorly characterised auxiliary protein, the agnoprotein (Chong et al. 2019).

Agnoprotein is a small, highly basic phosphoprotein found in a minority of polyomavirus. BKPyV and JC polyomavirus (JCPyV) are the only hPyVs reported to express agnoproteins. The agnoproteins of BKPyV and JCPyV are highly conserved in their amino terminus with an 83% sequence identity, suggesting a possible conserved function. Though, there have been multiple studies focussing on JCPyV agnoprotein, its exact function in the polyomavirus lifecycle remains unclear. Characterisation of agnoprotein in JCPyV and BKPyV has shown its subcellular localisation to be cytoplasmic, perinuclear, and within cellular membranes, which suggests a multipurpose function during infection (Gerits & Moens, 2012). The loss of agnoprotein expression leads to an egress defect in BKPyV infections, where infectious progeny virions accumulated in the nucleus (Panou et al. 2018). This accounts for the reduction in virus titres that have been observed in BKPyV infections when the agnoprotein was absent. However, a precise mechanism behind how the agnoprotein drives viral egress is lacking. Multiple binding partners have been described for JCPyV agnoprotein, whilst only a handful have been observed for BKPyV agnoprotein (Gerits & Moens, 2012). These interactions between agnoprotein and host factors have been validated, but there is limited information on how these virus-host interactions may relate to agnoprotein function during viral egress.

BKPyV and JCPyV agnoproteins share characteristics with a family of virally encoded proteins, termed viroporins. Viroporins are small pore forming proteins, containing hydrophobic regions that form at least one amphipathic helix which span cellular membranes (Nieva et al. 2012; Scott & Griffin, 2015; Royle et al. 2015). Viroporin-mediated membrane permeablisation functions in many different ways during viral life cycles. Influenza M2 protein functions as a proton channel during Influenza’s entry and egress in order to allow correct endosomal fusion and maturation of virions (Takeuchi & Lamb, 1994; Shimbo et al. 1996). Coxsackie virus B 2B protein modulates ER permeability to manipulate intracellular Ca^2+^ levels in order to allow for viral replication and release (van Kuppeveld et al. 1997). JCPyV agnoprotein has been shown to impair membrane integrity and homo-oligomerise to modulate intracellular Ca^2+^ levels, suggesting it functions as a viroporin (Suzuki et al. 2010; Suzuki et al. 2013). Here we have described a conserved viroporin function for BKPyV agnoprotein both *in vitro* and *in vivo*. Furthermore, we have identified small molecule inhibitors which inhibit BKPyV agnoprotein function *in vitro* and within the full BKPyV lifecycle, proving the possibility of developing antiviral compounds that target viral release through inhibition of agnoprotein.

## Methods and Materials

### Plasmids/Primers

pGEM7-BKPyV (strain Dunlop) plasmid (from Michael Imperiale, University of Michigan), was used to generate an agnoprotein knockout mutant via site-directed mutagenesis (Panou et al. 2018). GFP Coxsackie virus B (CoxV B) 2B and a viroporin mutant were kindly provided by Frank van Kuppeveld (University of Utrecht). For bacterial expression the BKPyV agnoprotein DNA sequence was codon optimised for E. coli (GeneWiz), and gene fragment amplified using: 5’ - AAA AAA GGT ACC ATG GTT CTG CGC CAG CTG AG - 3’, and 5’ - AAA AAA GGA TCC TTA GCT ATC CTT CAC GCT ATC TTT CAC G - 3’. The PCR fragment was cloned into pET19b between Kpn1 and BamHI. GFP BK Agno was cloned into peGFP-C1 between EcoR1 and BamHI, using the following primers: 5’ ATA TAT GAA TTC CAT GGT TCT GCG CCA GCT GTC 3’; 5’ ATA TAT GGA TCC CTA GGA GTC TTT TAC AGA G 5’; 5’ ATA TAT GAA TTC CAT GGT TCT TCG CCA GCT G 3’; and 5’ ATA TAT GGA TCC CTA TGT AGC TTT TGG TTC AGG C 3’.

### Cell Culture

HEK 293 cells were maintained at 37 °C, 5 % CO2 in Dulbecco’s minimum essential media (DMEM), supplemented with 10 % foetal bovine serum (FBS) and 1 % penicillin/streptomycin (P/S). HEK 293 cells were transfected when required using PEI at a ratio of 1:4 (DNA: PEI) in OptiMEM. Renal proximal tubular epithelial (RPTEs) cells were maintained at 37 °C, 5 % CO2 in renal epithelial growth media with the REGM Bulletkit supplements and passaged no higher than passage 7, as described (Panou et al. 2020).

### Cell infection, drug treatment, and quantification of viral release

RPTE cells were infected with BKPyV for 2hr at 37 °C, before virus was removed and fresh media applied. Drugs were added to infected cells 24 hr post-infection in fresh media, at 48 hr post-infection media was harvested. Media was applied to fresh RPTEs for 2 hr, and then replaced with fresh media. RPTE cells were incubated for 48hr prior to paraformaldehyde fixation. Cells were then permeabilised with 0.1% Triton x100 and blocked in 5% BSA, before being stained for VP1 using αVP1 (pAb597). Analysis of VP1 positive cells was carried out the IncuCyte ZOOM instrument (Essen BioScience, Ann Arbor, MI, USA). The software parameters with a 10× objective were used for imaging.

### Cell Viability Assay

RPTE cells were incubated with drugs for 24 hr. After incubation media was replaced with 1 mg/ml MMT reagent in OptiMEM and placed at 37 °C in the dark for 30 minutes. MTT reagent was then removed, and DMSO was used to resolubilise precipitate that had formed. Samples were analysed via absorbance at 560 nm.

### Expression of recombinant 10xHis BKPyV agnoprotein

pET19b BKPyV agnoprotein was transformed into DE3 gold cells, and used to inoculate 1 L of LB broth. Bacterial cultures were grown to 0.6 OD600, then induced with 1 mM IPTG. After induction, bacterial cultures were incubated overnight at 37 °C on an orbital shaker. Bacterial cultures were harvested via centrifugation at 5000 RMP, and pelleted cells lysed in 50 ml of lysis buffer (10 mM Tris pH8, 500 mM NaCl, 1% Triton x100, protease inhibitor cocktail, 50 mg/μl lysozyme, 1 μl Benzonase). Bacterial lysate was clarified via centrifugation at 4000 RMP, and soluble fraction applied to NiNTA resin. NiNTA resin was washed in 3 bed volume (bv) of 10 mM Tris pH8, 500 mM NaCl, 25 mM imidazole. Protein was then eluted from NiNTA using increasing concentrations of imidazole: 50, 100, 200, 400 mM; 2 bv of each concentration was applied to the resin. 3 bv of buffer containing 1 M imidazole was then passed through the column to elute any remaining protein. Fractions collected were analysed for protein on 15% SDS PAGE using Instant Blue staining. Fractions positive for 10xHis BKPyV agnoprotein were carried forward to ion exchange chromatography. Samples were diluted 10-fold to lower salt concentration, before being applied to HiTrap SP column, and protein eluted using a NaCl gradient from 50 mM to 1 M. Collected fractions were analysed for protein on 15% SDS PAGE using Instant Blue staining. Recombinant agnoprotein was concentrated using C4 solid phase extraction columns. Fractions from the ion exchange chromatography were applied to the columns and washed with 5% acetonitrile to remove the salt. Bound protein was then eluted in 95% acetonitrile, dried down under vacuum and dissolved in DMSO.

### GST pull down

Lysates from HEK 293 cells expressing GFP-agnoprotein and GST-agnoprotein were incubated overnight with glutathione beads. Precipitates were washed four times with lysis buffer and twice in TBS. Proteins were eluted from the beads by resuspension in SDS PAGE loading buffer prior to western blot analysis.

### Merocyanine 540 assay

HEK 293 cells transfected with GFP, GFP BKPyV agnoprotein and GFP CoxV B 2B proteins were disassociated in PBS-based enzyme-free disassociation buffer 48 hr post-transfection. Cells were collected via a gentle centrifugation and resuspended in 1ug/ml Merocyanine 540. Cells were incubated at 37 °C for 10 mins, and then collected via centrifugation. Cells were resuspended in PBS and analysed on Beckman Fortessa at 488 nm and 620 nm.

### Cytoplasm/Membrane subcellular fractionations

BKPyV infected RPTE cells were harvested at 72 hrs. Cells were resuspended in M1 (10 mM PIPES pH 7.4, 0.5 mM MgCl2, Protease inhibitors) and sonicated for three 30 second bursts. Cell lysate was adjusted with M2 (10 mM PIPES pH 7.4, 600 mM KCl, 150 mM NaCl, 22.5 mM MgCl2) at a ratio of 1:4 (M2:M1). Centrifugation was carried out at 4 °C, 3000 xg for 10 minutes to pellet nuclei and intact cells. Supernatant was collected and ultracentrifugation at 100,000 xg for 30 mins was performed. Resulting supernatant (cytoplasmic fraction) was precipitated with 4 volumes of acetone, and pellet (membrane fraction) was washed 3 times with adjusted M1, and once with 70% ethanol. Cytoplasmic and membrane pellets were resuspended in SDS loading buffer for analysis. Alternatively, membrane pellets were resuspended in adjusted M1 containing either 1% Triton X100, 1 M KCl, or 1 M NaOH and incubated at room temperature for 30 mins. Resuspended membrane pellets were ultracentrifuged at 100,000 xg for 10 mins, and supernatant analysed by western blotting.

### Preparation of unilamellar liposomes

L-α -phosphatidic acid (egg monosodium salt) (PA) and L-α -phosphatidylcholine acid (egg) (PC) solubilised in chloroform were mixed 1:1 with a final concentration of 0.5% (wt. /wt.) L-α-phosphatidylethanolamine with lissamine. Lipid mixture was dried under a stream of argon and placed in a vacuum for 2 hr at room temperature. Dried lipids were rehydrated at room temperature overnight in a self-quenching concentration of carboxyfluorescein (CF) (Sigma-Aldrich) buffer (50 mM CF in HEPES-buffered saline [HBS] [10 mM HEPES-NaOH (pH 7.4), 107 mM NaCl]).

Unilamellar liposomes were then formed at 37 °C via extrusion through a 0.4 μm Nuclepore Track-Etch membrane filter (Whatman), using an Avanti miniextruder with Hamilton glass syringes. Liposomes produced were washed three times and purified by ultracentrifugation to remove free dye. After purification liposomes were resuspended in HBS and quantified via absorbance at 520 nm.

### Liposome permeability assay

Release of liposome content was monitor by fluorimetry, utilising the self-quenching properties of carboxyfluorescein. Fluorescence measurements were performed with a FLUOstar Optima microplate reader (BMGTechnologies) with excitation and emission filters set to 485 nm and 520 nm, respectively. 50 μM of liposomes were incubated with either 1% Triton, recombinant 10xHis BK agnoprotein, or 1 μM melittin, and fluorescence measurements were started immediately every 30 secs for a total of 30 mins. For inhibitor studies, recombinant 10x His BK agnoprotein was pre-incubated with each compound for 10 minutes at room temperature prior to the start of the assay

## Results

### BKPyV agnoprotein self-associates and is associated with membranes in kidney cells

The agnoproteins of BKPyV and JCPyV are highly conserved and comparatively across the full peptide sequence they share 80% similarity. Both proteins share a hydrophobic region of ~21 amino acids in the middle of the protein sequence (Fig 1A–1B), with high helical tendency (Fig 1B). NMR studies using synthesised JCPyV agnoprotein (Coric et al. 2014) have confirmed the existence of a stable helical structure between residues 24-39. This predicted helical domain is amphipathic with two hydrophobic sides, an aromatic side and a hydrophilic side, which suggests this helical domain may function both as a transmembrane domain and potential dimerization surface within the protein. Both of these are essential features for viroporin function. Crucially, JCPyV agnoprotein self-association has been confirmed in cells (Suzuki et al. 2010). In contrast, it is less clear if BKPyV can self-associate in cells. To address this, HEK293 cells were co-transfected with plasmids expressing GST-agnoprotein and GFP-agnoprotein and the GST fusion protein captured on glutathione agarose beads. GFP-agnoprotein was unable to be pulled down by GST alone but bound efficiently to GST-agnoprotein (Fig 1C).

**Figure 1.**
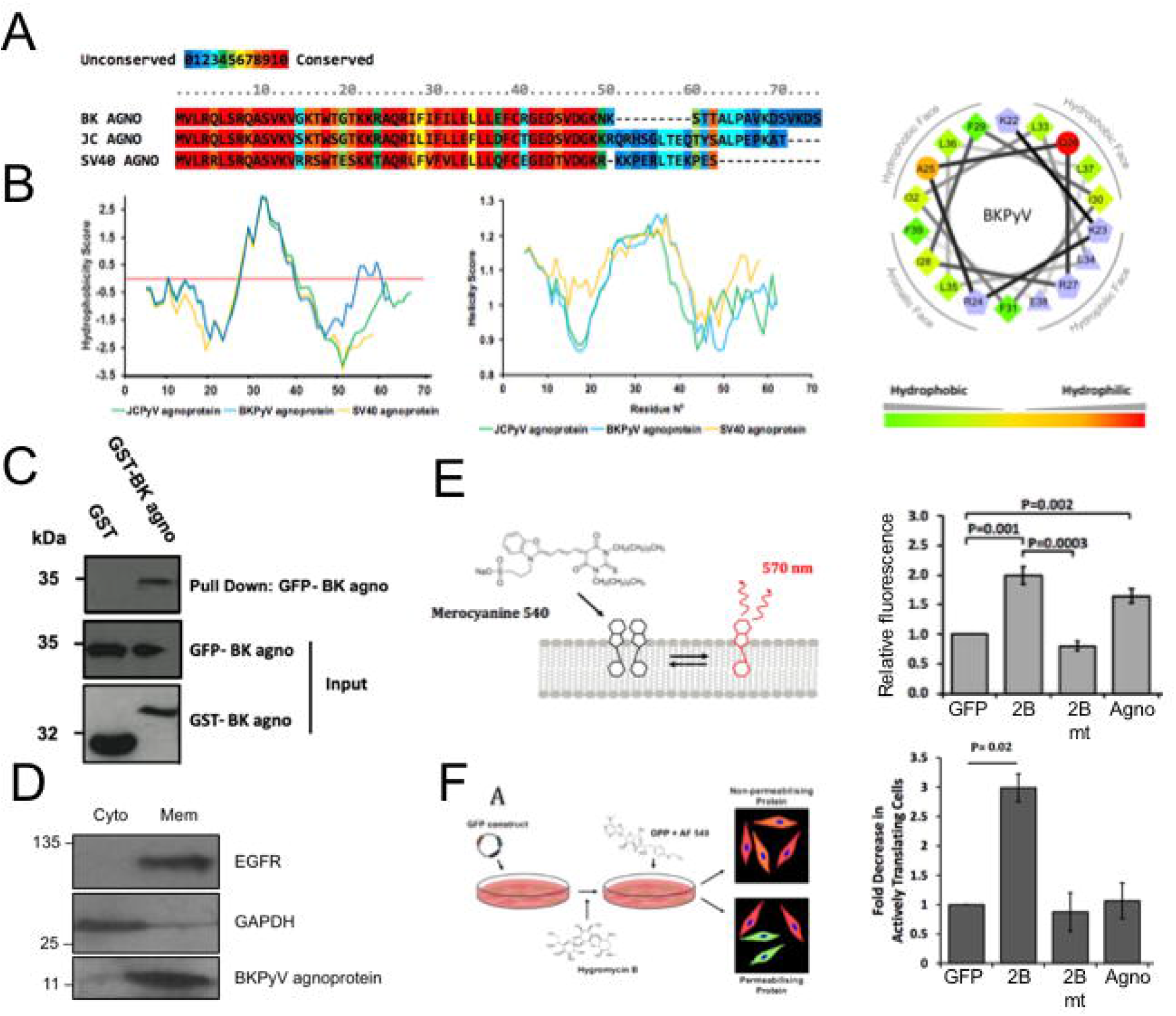
BKPyV agnoprotein displays the properties of a viroporin. (A) Sequence alignment of BKPyV, JCPyV and SV40 agnoproteins, performed by PRALINE server. (B) Hydrophobicity and helicity scores across the BKPyV, JCPyV and SV40 agnoproteins. A helical wheel diagram representation of the central helical structure of BKPyV agnoprotein with each amino acid coloured either on the displayed hydrophobic to hydrophilic scale (green to red) or blue for polar residues. (C) Western blot analysis of GST pulldown using GST or GST-BKPyV agnoprotein to pulldown lysates from cells expressing GFP-BKPyV agnoprotein. Pulldowns are probed for antibodies against GST (input) and GFP (input and pulldown). N=3. (D) Western blot analysis of membrane/cytosol fractionation from RPTE cells infected with BKPyV (MOI 1). Fractions were probed with antibodies against EGFR (membrane), GAPDH (cytosol) and agnoprotein. N=3. (E) Schematic diagram showing the structure of MC540 and the equilibrium formed in cellular membranes. Histogram quantifying the MC540 fluorescent positive population of cells relative to GFP and GFP fusion proteins. N=4. (F) Schematic of the workflow for analysis of plasma membrane integrity using hygromycin and OPP. Histogram quantifying the fold decrease in translationally active cells expressing GFP and GFP fusion proteins after hygromycin treatment. N=3. Data show mean values with SD, analysed with a two-tailed unpaired t-test. Significance is highlighted on the graph.

Next, we attempted to further define the membrane interaction properties of BKPyV agnoprotein. Analysis of sub-cellular fractions of RPTE cells infected with BKPyV showed that agnoprotein was predominantly localised in the membrane compared with the cytosolic fraction (Fig 1D). To examine if agnoprotein disrupts cellular membranes, we examined cellular membrane structures using Merocyanine 540 (MC540), a fluorescent dye that can measure the effect of lipid packing in membranes and has been used extensively to probe cellular membrane structure, function and integrity (Mandall et al. 1999; Verkman et al. 1987; Williamson et al. 1983). We used flow cytometry to examine MC540 fluorescence of HEK293 cells transfected with GFP-agnoprotein compared to GFP alone. As a positive control, we made use of a GFP coxsackie B virus 2B protein, an extensively characterised viroporin known to cause membrane perturbation, and a non-functional 2B mutant (L46N/V47N/I49N/I50N) impaired in oligomerisation and pore forming activity (de Jong et al. 2004). The MC540 intensity in agnoprotein and 2B expressing cells was significantly higher than in either the GFP or 2B mutant expressing cells (Fig 1E), suggesting that agnoprotein disrupts lipid packing and modifies the structure of membranes. To extend these findings, we performed a Hygromycin B permeability assay. Hygromycin B is an inhibitor of translation and intact mammalian cells are impermeable to hygromycin B. However, hygromycin B can enter cells where the plasma membrane has been permeabilised. HEK 293 cells were transfected with GFP-BKPyV agnoprotein, GFP-2B (wildtype and mutant) and empty GFP plasmid. Following incubation for 48 hours, cells were treated with low concentrations of hygromycin B and O-propargyl-puromycin, and copper(I)-catalysed azide alkyne cycloaddition of a functionalised fluorescent probe was used to enable visualisation of active protein translation. In cells transfected with GFP-2B, levels of fluorescence were significantly decreased as anticipated given previous reports of the membrane permeabilising effects of this viroporin. In contrast, in agnoprotein transfected cells, levels of translation driven fluorescence were unaffected by hygromycin B and remained at similar levels to the 2B viroporin mutant and GFP alone (Fig 1F). Thus, whilst agnoprotein alters cellular membranes it does not enhance plasma membrane permeability.

### Generation of high-purity recombinant BKPyV agnoprotein from bacteria

The characterisation of agnoprotein has been hampered by the lack of an efficient system to purify recombinant proteins compatible with down-stream biochemical analyses. Here we have improved upon a method used previously to purify the human papillomavirus (HPV) E5 protein (Wetherill et al. 2012; Wetherill et al. 2018) to produce a ‘detergent free’ recombinant agnoprotein for analysis in liposomal membranes. In our modified protocol, a 10-His epitope is fused to the amino terminus of agnoprotein. A significant portion of this fusion protein can be solubilised in Triton-X100 and purified by nickel chromatography with an imidazole gradient (Fig 2A-B). Purity is increased when fractions are taken forward for cation exchange chromatography (cIEX) and agnoprotein eluted with a NaCl gradient. Figure 2C shows western analysis from the final concentration step, including the clear presence of higher order oligomers. When further analysed by SDS PAGE, recombinant agnoprotein displayed characteristic SDS-resistant dimeric and oligomeric species (Figure 2D). This distinct pattern of oligomerisation was concentration dependent, with tetramers and pentamers clearly visible at higher concentrations.

**Figure 2.**
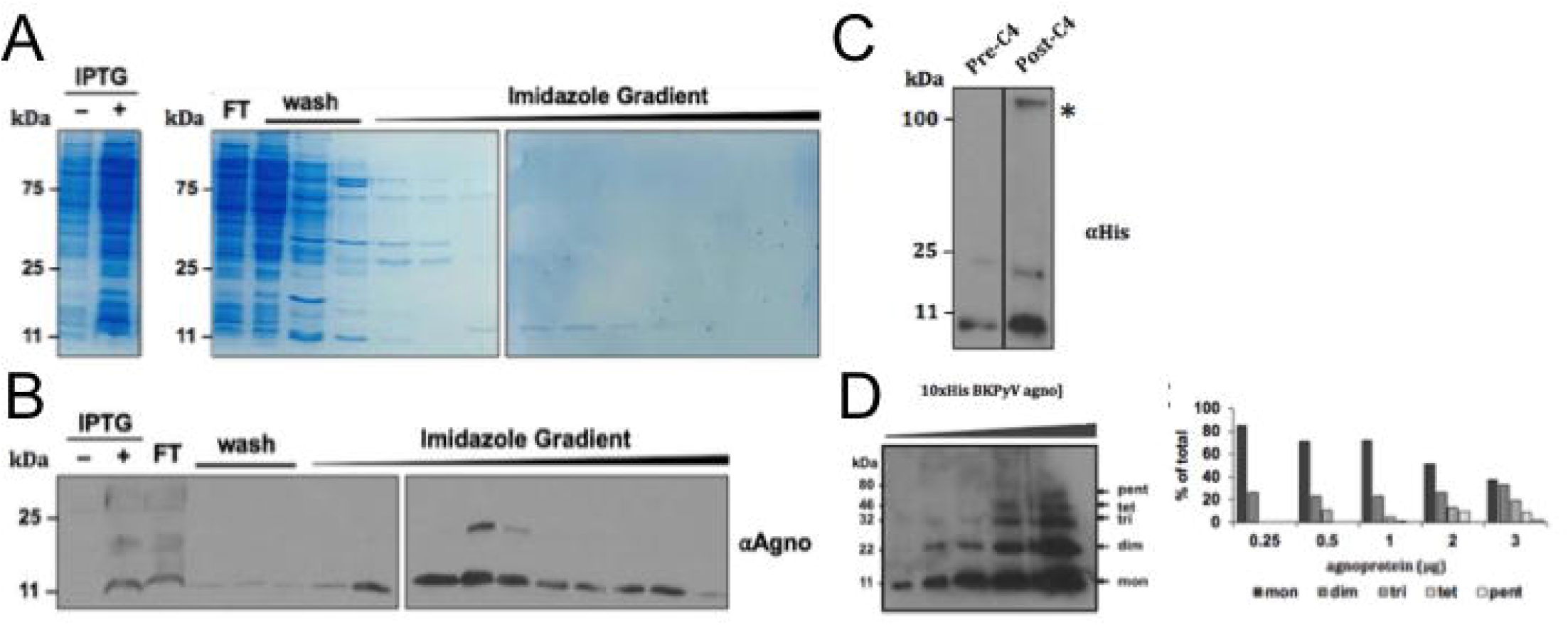
Production of recombinant BKPyV agnoprotein. (A) SDS PAGE of IPTG induced His-BKPyV agnoprotein fractions collected during Ni-affinity purification, stained with Coomassie blue. (B) Western blot of fractions collected during Ni-affinity purification, probed with anti-agnoprotein antibody. (C) Western blot analysis pre-C4 and post-C4 concentration steps, probed with anti-His antibody. (D) SDS PAGE of increasing concentrations of His-BKPyV agnoprotein, probed with anti-agnoprotein antibody. Densitometry analysis of the oligomeric species observed on the SDS PAGE gel.

### Recombinant BKPyV agnoprotein associates with liposomes

To determine whether recombinant agnoprotein associated with membranes, protein-liposome suspensions were subjected to ultracentrifugation, resulting in flotation through a discontinuous Ficoll gradient, as previously described (Wetherill et al. 2012). Gradient fractions were analysed by western blotting to detect agnoprotein and the distribution of liposomes was assessed by rhodamine fluorescence. This confirmed that the liposomes had migrated to the 10% Ficoll-aqueous buffer interface (Figure 3A) and that the majority of the agnoprotein associated with the liposomes (Figure 3B top blot). Treatment with Triton-X100 detergent disrupted the liposome-agnoprotein interaction (Figure 3B bottom blot), indicating that the migration of the protein to the Ficoll-aqueous buffer was liposome-dependent.

**Figure 3.**
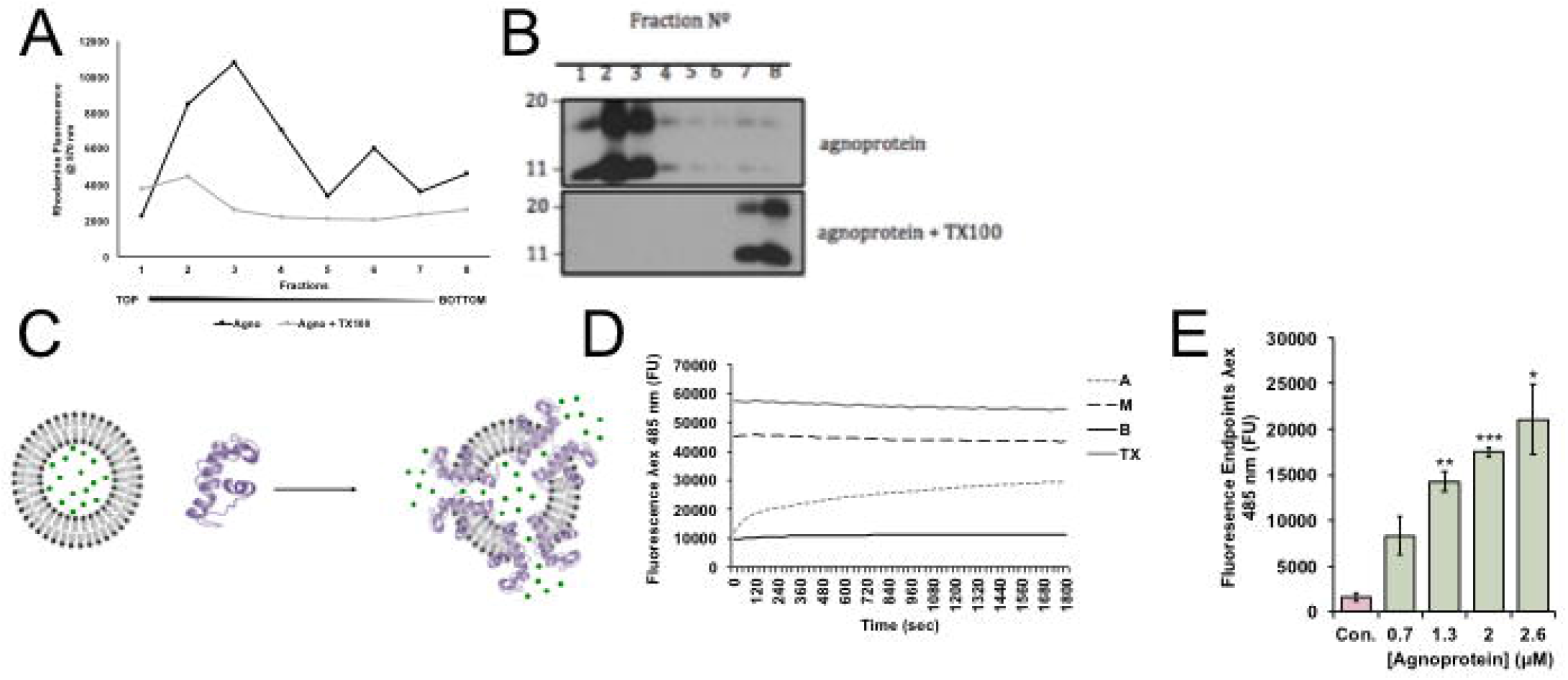
BKPyV agnoprotein displays channel forming activity. (A) Rhodamine fluorescence measured across fractions taken from Ficoll gradients to track the migration of liposomes. (B) Western blot analysis of fractions taken from Ficoll gradients to monitor the migration of the agnoprotein, probed with an anti-agnoprotein antibody. (C) Schematic of the dye-release assay. (D) Carboxyfluorescein release over the course of 30 minutes after liposome incubation with Triton-X100 (T), melittin (M) or His-BKPyV agnoprotein (A). Dye release measured by relative end point fluorescence of increasing concentrations of His-BKPyV agnoprotein. All experiments performed minimum of N=3. Data show mean values with SD, analysed with a two-tailed unpaired t-test. Significance is highlighted on the graph.

### Recombinant agnoprotein shows channel activity in liposomes

To address the functional implications of the agnoprotein membrane association, we employed a well-utilised liposome-based fluorescent dye release assay used previously to investigate viroporin function (Wetherill et al. 2012; St-Gelais et al. 2007) (Figure 3C). Increasing amounts of agnoprotein were incubated with liposomes containing the fluorescent dye carboxyfluorescein (CF) at self-quenching concentrations. The release of this dye resulted in the recovery of fluorescent signal, which was detected in real time by a fluorimeter. The pore forming component of bee venom (Melittin – M) was used as a positive control and treatment with the detergent Triton-X100 (TX) resulted in maximum fluorescence (Figure 3D–3E). Baseline readings were calculated from solvent controls (10% DMSO and liposomes - B). The addition of agnoprotein (A) promoted a rapid release of CF from liposomes (Figure 3D).

### Agnoprotein viroporin activity shows differential sensitivity to classical viroporin inhibitor compounds

Several prototypic classes of inhibitor compounds have been shown to abrogate viroporin function in vitro, including the adamantanes, rimantadine and amantadine, and nonylated imino sugars (e.g. *N*N-DNJ) (Scott and Griffin 2015). These compounds have since been shown to exert antiviral effects against a number of viruses including HCV, BVDV, Dengue and HPV, (StGelais 2009, Scott & Griffin, 2015; Wetherill 2012; Wetherill 2018). Despite this, viroporin inhibitors have never been tested against any agnoprotein. Incubation of agnoprotein with high concentrations of amantadine (400 uM) did not significantly affect the release of CF from liposomes, as measured by endpoint fluorescence (Figure 4A). However, the same concentration of rimantadine reduced channel activity by approximately 50% (Figure 4A). Addition of *N*N-DNJ also led to a significant reduction in agnoprotein mediated CF release from liposomes (Figure 4A). 3,3’-Diisothiocyano-2,2’-stilbenedisulfonic acid (DIDs) has been shown to inhibit BKPyV release (Evans et al. 2015). Whilst the mechanism by which DIDs inhibits virus release remains unknown, this compound has been shown to inhibit Enterovirus 2B viroporin activity. Treatment of agnoprotein with DIDs led to an ~80% reduction in CF release (Figure 4A).

**Figure 4.**
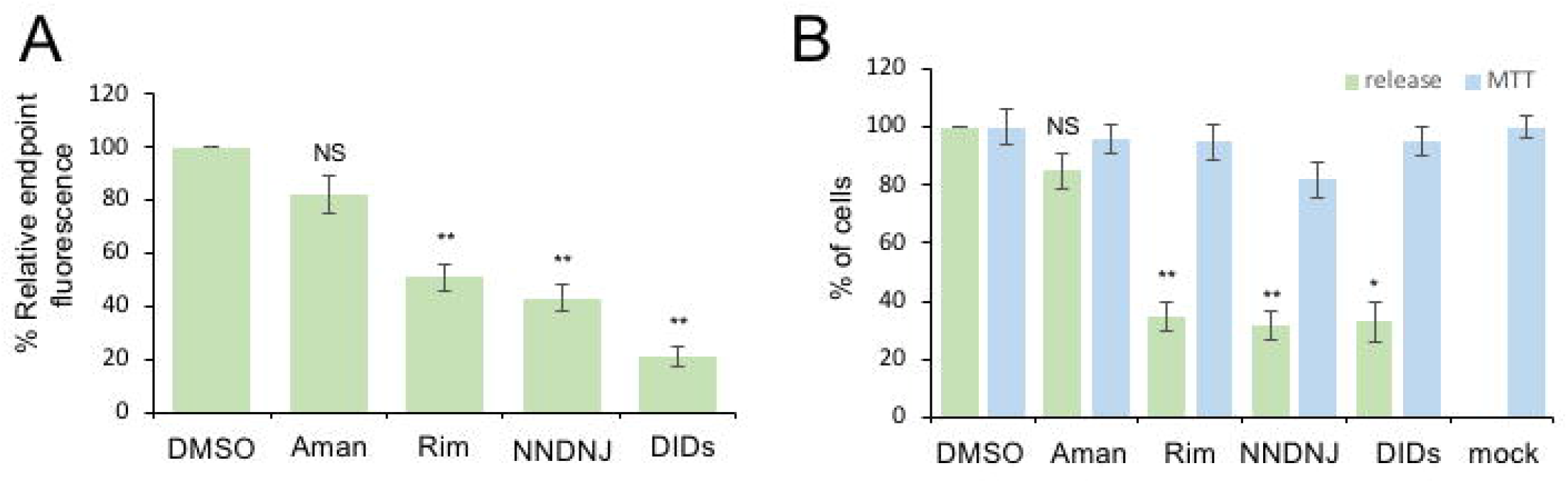
Prototypic viroporin inhibitors inhibit BKPyV agnoprotein mediated dye release and reduce BKPyV release from infected RPTE cells. (A) Endpoint fluorescence of liposomes incubated with His-BKPyV agnoprotein and viroporin inhibitors relative to a DMSO control. (B) Quantification of VP1 positive RPTE cells using an IncuCyte ZOOM relative to a DMSO control. Cell viability assays of RPTE cells treated with viroporin inhibitors. All experiments performed minimum of N=3. Data show mean values with SD, analysed with a two-tailed unpaired t-test (*P<0.05, **P>0.01).

### Viroporin inhibitors reduce BKPyV release from infected kidney cells

To further investigate if viroporin function contributes to the essential role of agnoprotein in mediating BKPyV release from infected cells, RPTE cells were infected with 0.5 MOI BKPyV and compounds added in fresh media at 24 hours post infection. Media from treated and control cells was harvested at 48 hours post infection, at a point that we have previously detected released virus (Panou et al. 2018) and applied to naïve RPTE cells to enable us to quantify the level of virus released. A significant reduction in released virus was observed for rimantadine, NN-*D*NJ and DIDs (Figure 4B). Similar to the results of the dye release assay, Amantadine treated cells showed no significant reduction in virus release compared to solvent control (Figure 4B). Crucially, at the concentrations tested the inhibitor compounds had no significant impact on cell viability (Figure 4B).

## Discussion

The precise molecular mechanisms by which agnoprotein contributes to BKPyV release from an infected cell remain elusive. In this study we provide evidence for a previously undocumented function of BKPyV agnoprotein as a viroporin. Viroporins have been shown to play essential roles in virus lifecycles across many viral families. These roles are usually associated with viral entry and release, where with many viruses it is important that the local concentration of ions is carefully rebalanced to prevent aberrant viral disassembly (Scott et al. 2015). Using cell biology assays, the agnoprotein of JCPyV has been proposed as a potential viroporin (Suzuki et al. 2015). Agnoproteins are found in many members of the *Polyomaviridae* and are highly conserved. It is unknown if viroporin function was conserved across the members, and the role of this function is unknown in the context of the viral lifecycle [Gerits et al. 2012].

Key to our discovery was the establishment of the first robust system for the expression and purification of a recombinant BKPyV agnoprotein, which should permit future comprehensive biophysical and structural characterisation of the agnoprotein. Our initial studies confirmed that recombinant agnoprotein exists as an oligomer in membrane like environments. As seen previously for HPV E5 and HCV p7 (Wetherill et al. 2012; St-Gelais et al. 2007), SDS acts both as a membrane mimetic and as a denaturant, leading to a laddering effect of agnoprotein oligomeric species by SDS PAGE, with higher-ordered forms being less abundant than monomers or dimers.

Viroporins are attractive targets for antiviral therapy with the adamantane compounds clinically available targeting Influenza A M2 protein (Hay et al. 1985). Our study finds that agnoprotein is resistant to high concentrations of amantadine but can be inhibited by relatively high concentrations of rimantadine. This highlights differences between agnoprotein and the prototypic viroporins M2 and p7, several variants of which can be highly sensitive to both compounds. We also demonstrated that both the imino sugar *N*N-DNJ and DIDs reduced agnoprotein channel activity significantly better than adamantanes *in vitro*. However, whilst lacking true drug like potency, these prototypic viroporin inhibitors can be useful for identifying both potential binding sites and inhibitory modes of action that can subsequently be targeted via rational design or compound screening approaches.

A major aspect of agnoprotein function is to aid in virus release from infected cells. Our study suggests that viroporin activity is necessary for this function as treatment of infected cells with several viroporin inhibitors resulted in reduced virus release into the media. Our data aligns with a recent published observation identifying DIDs as an inhibitor of BKPyV release (Evans et al. 2015). Data from agnoprotein knockout models indicates that agnoprotein is necessary for the efficient egress of infectious BKPyV virions from the nucleus into the cytoplasm. How this is achieved in a viroporin-dependent mechanism is not yet understood. It is unlikely that agnoprotein channels would directly mediate transport of BKPyV virions given the potential size of such channels and so it is more likely that the viroporin activity is used to manipulate the host secretory system and trafficking pathways from the nucleus to the cytoplasm.

In summary, our data show a new function for the BKPyV agnoprotein as a viroporin necessary for virus release. Our findings provide further evidence for a virus regulated release mechanism.

## Acknowledgements

We thank Michael Imperiale (University of Michigan), Denise Galloway (Fred Hutchinson Cancer Research Center), Frank van Kuppeveld (Utrecht University), Ugo Moens (Tromso University Norway) and Chris Buck (National Cancer Institute) for providing essential reagents and advice.

